# Predicting humoral alloimmunity from differences in donor-recipient HLA surface electrostatic potential

**DOI:** 10.1101/294066

**Authors:** Dermot H Mallon, Christiane Kling, Matthew Robb, Eva Ellinghaus, J Andrew Bradley, Craig J Taylor, Dieter Kabelitz, Vasilis Kosmoliaptsis

## Abstract

In transplantation, development of humoral alloimmunity against donor HLA is a major cause of organ transplant failure but our ability to assess the immunological risk associated with a potential donor-recipient HLA combination is limited. We hypothesised that the capacity of donor HLA to induce a specific alloantibody response depends on their structural and physicochemical dissimilarity compared to recipient HLA. To test this hypothesis, we first developed a novel computational scoring system that enables quantitative assessment of surface electrostatic potential differences between donor and recipient HLA molecules at the tertiary structure level (electrostatic mismatch score-three dimensional; EMS-3D). We then examined humoral alloimmune responses in healthy females subjected to a standardised injection of donor lymphocytes from their male partner. This analysis showed a strong association between the EMS-3D of donor HLA and donor-specific alloantibody development; this relationship was strongest for HLA-DQ alloantigens. In the clinical transplantation setting, the immunogenic potential of HLA-DRB1 and -DQ mismatches expressed on donor kidneys, as assessed by their EMS-3D, was an independent predictor of development of donor-specific alloantibody after graft failure. Collectively, these findings demonstrate the translational potential of our approach to improve immunological risk assessment and to decrease the burden of humoral alloimmunity in organ transplantation.

## Introduction

The Human Leukocyte Antigen (HLA) gene complex encodes highly polymorphic proteins that are the main immunological barrier to successful cell, tissue and organ transplantation. Immune recognition of HLA class I and class II expressed on donor tissue stimulate the development of donor-specific antibodies (DSA) that are the major cause of organ transplant failure in the medium- to long-term (1-5). Moreover, development of alloantibody through pregnancy, blood transfusion and previous transplantation may severely limit the opportunity for organ transplantation (6, 7). Current strategies to offset the risk for development of DSA and of antibody-mediated rejection focus on minimising the number of HLA mismatches between donor and recipient and on the administration of immunosuppression regimens that aim to suppress the recipient immune response. HLA matching is incorporated into many deceased donor organ allocation schemes, but because of the extensive polymorphism of the HLA system and the relative limitation in the size of the donor organ pool, most recipients receive allografts with one or more mismatched HLA alleles. HLA incompatible allografts necessitate the use of increased immunosuppression and this is a major cause of recipient morbidity and mortality (8, 9).

Current assessment of the immunological risk associated with a particular transplant is based on enumerating the number of HLA mismatches between donor and recipient and is predicated on the assumption that all mismatches within an HLA locus are of equal significance to graft outcomes. However, it is clear from animal studies that humoral alloimmunity is critically dependent on the nature of the Major Histocompatibility Complex (MHC) mismatch between donor and recipient and this has been supported by observational studies in humans suggesting that certain donor HLA are tolerated by the recipient immune system (10, 11). Recent evidence shows that the potential of donor HLA to induce humoral alloresponses (HLA immunogenicity) might be a function of the number and location of amino acid sequence polymorphisms compared to recipient HLA molecules (12, 13). Numerous studies support an association between donor HLA immunogenicity, considered at the amino acid sequence level, and the likelihood of DSA developing after transplantation and that this approach might offer superior assessment of donor-recipient histocompatibility compared to conventional HLA matching strategies (14-17). Studies by our group have shown that the predictive ability of sequence-based HLA immunogenicity algorithms can be significantly enhanced by consideration of the physicochemical properties of amino acid polymorphisms expressed on donor HLA molecules (18-21).

Despite its promise, sequence-based assessment of HLA immunogenicity does not account for the conformational nature of antigenic recognition by B cell receptors and for the effect of individual amino acid polymorphisms on B cell epitope structure and physicochemical properties (e.g. surface exposure, polarity, surface charge, hydrophobicity) (22-24). In particular, antibody-antigen interactions are largely governed by electrostatic forces dictated by the number and distribution of charged atoms on the surface of the HLA molecule (24-27). We have previously shown that, despite variation in their amino acid composition, HLA B cell epitopes are characterised by unique surface electrostatic potential properties that explain serological patterns of HLA-specific antibody binding (28-30). In this study, we hypothesised that the capacity of donor HLA to induce a specific alloantibody response can be predicted by quantitative assessment of their structural and surface electrostatic potential differences compared to recipient HLA molecules. We have developed a novel computational scoring system to quantify and compare HLA electrostatic properties that utilises molecular modelling techniques, structural information from X-ray crystallography, and application of protein electrostatics theory. This approach was validated by analysis of HLA-specific antibody responses in a unique model of HLA sensitisation comprising patients that underwent a single injection of donor lymphocytes (lymphocyte immunotherapy) in a defined donor-recipient setting without the influence of immunosuppression or interference from other sensitising events. The applicability of our findings in transplantation was then examined by analysis of DSA responses in patients listed for repeat renal transplantation.

## Results

### Computational approach for calculation of HLA surface electrostatic potential and quantification of differences in electrostatic potential between HLA molecules

We generated a bioinformatics protocol to enable HLA structure prediction, surface electrostatic potential calculation and quantification of electrostatic potential differences between different HLA molecules (Figure 1). Because the structure of very few HLA molecules has been determined experimentally, the atomic resolution structure of a given HLA class I or class II molecule was calculated using comparative structure modelling, based on information derived from high quality HLA structures resolved by X-ray crystallography (using the program Modeller). Following validation of model structural quality, HLA molecules were computationally immersed in an aqueous solvent and the electrostatic potential in three-dimensional (3D) space surrounding the HLA structure was calculated numerically by solving the linearised Poisson-Boltzmann equation (as implemented in the program APBS). After structure superimposition, comparisons of electrostatic potential between two HLA molecules of interest were performed in a defined region of space above the HLA molecular surface and values expressed as electrostatic similarity distance (ESD), based on the Hodgkin index (as detailed in methods).

**Figure 1.**
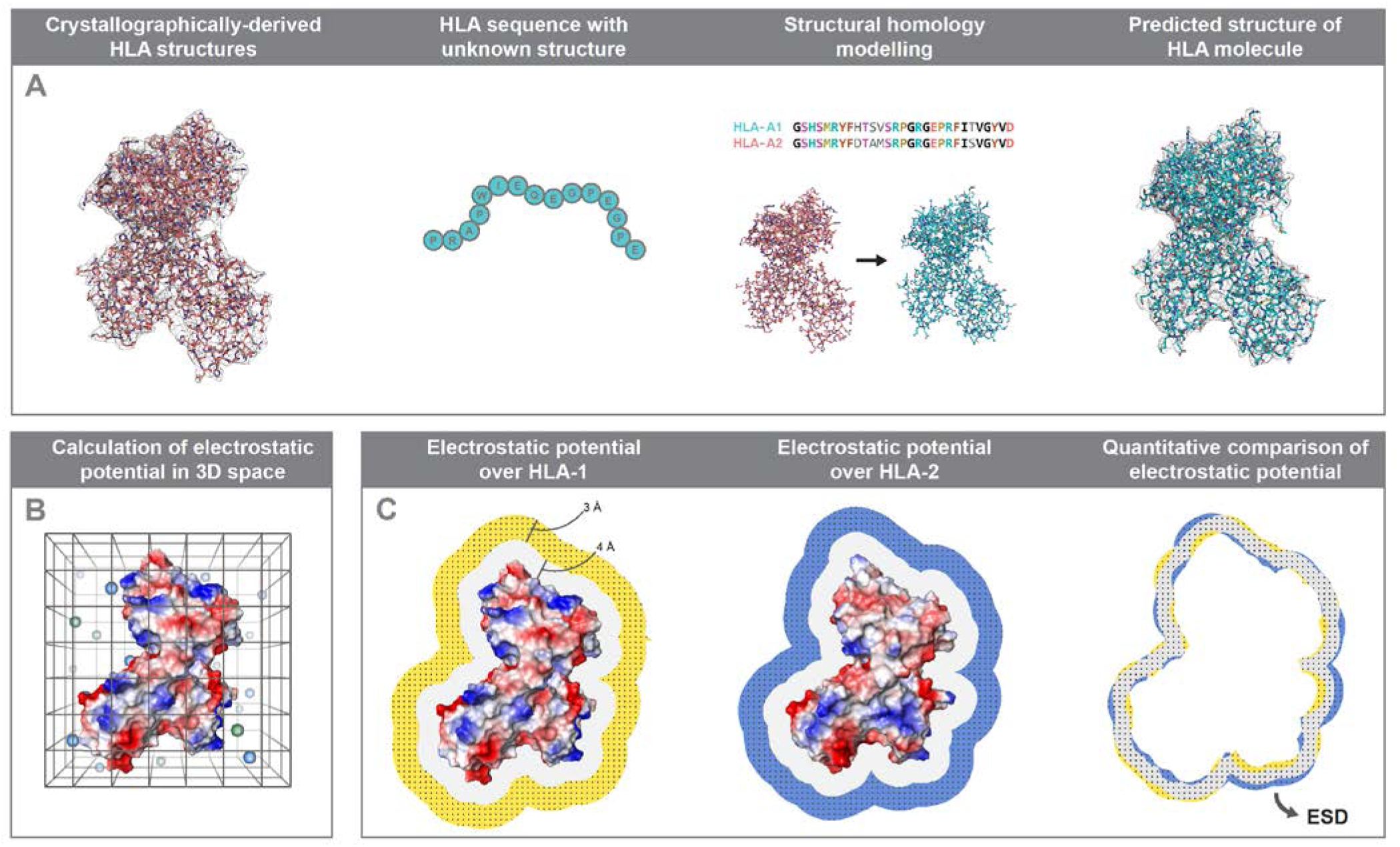
Schematic overview of the computational approach for quantification of surface electrostatic potential differences between HLA molecules. Bioinformatics approach to enable HLA structure prediction, surface electrostatic potential calculation and quantification of electrostatic potential differences between two HLA molecules. Panel A: The atomic resolution structure of a given HLA class I or class II molecule is calculated using homology modelling (Modeller) based on information derived from high quality HLA structures resolved by X-ray crystallography. Panel B: The electrostatic potential in three-dimensional (3D) space surrounding an HLA structure is calculated numerically by solving the linearised Poisson-Boltzmann equation for each point on a cubic grid (spacing of 0.33 Å, solvent ionic strength of 0.15 M, pH: 7.4). Panel C: Electrostatic potential comparisons consider cubic grid points within a defined region or ‘layer’ of space (of thickness δ=3Å) at a distance σ (4Å) above the van der Waals surface of the HLA molecule. Quantitative comparison of the electrostatic potential between two HLA molecules of interest are performed using the Hodgkin similarity index for grid points within the intersection of their ‘layers’ (depicted in grey), after the two structures are superimposed, and values are converted into a distance [Electrostatic Similarity Distance (ESD)].

The overall electrostatic potential disparity of a given donor HLA compared to recipient HLA molecules was quantified based on the three-dimensional electrostatic mismatch score (EMS-3D). As described in methods and shown in supplementary Figure S1, for HLA class I alloantigens, the mismatched donor HLA was compared electrostatically to each of the recipient HLA class I molecules to derive the respective ESDs and the minimum ESD value was taken to represent the EMS-3D (inter-locus comparison (21)). Similarly, for HLA class II alloantigens, the mismatched donor HLA was compared electrostatically to each of the recipient HLA within the same locus to derive the ESDs and the minimum value was taken to represent the EMS-3D (intra-locus comparison (19)).

### Amino acid sequence polymorphism and disparities in surface electrostatic potential among HLA class I and class II alleles

To investigate the relationship between amino acid sequence polymorphisms among HLA molecules and differences in their surface electrostatic potential we performed pair-wise, all-versus-all comparisons between common (frequency >1%) HLA alleles within individual HLA class I and class II loci. The ESD between HLA ranged from 0.00 to 0.777 (median: 0.307, IQR: 0.219-0.379), reflecting the overall structural and physicochemical similarity of molecules within the same protein family (Supplementary Table S1). Overall, there was poor correlation between amino acid sequence polymorphism and electrostatic disparity for compared HLA class I (R^2^=0.439) and class II molecules (R^2^=0.317) with wide variation of ESD values for the same level of sequence polymorphism (supplementary Figure S2). It was notable that disparities in electrostatic potential were highest among HLA-DQ alloantigens and lowest among HLA-DR and -DP alloantigens, whereas HLA class I molecules had similar levels of variation in their electrostatic properties. Supplementary Figure S3 shows the ESDs for pairs of compared HLA alleles within individual HLA loci presented as symmetrical heat maps with re-ordering such that electrostatically similar alleles are clustered together.

### Analysis of HLA-specific antibody responses after lymphocyte immunotherapy

Investigation of alloantibody responses against donor HLA in human transplantation is commonly confounded by many, often difficult to control, factors such as differences in allosensitisation events [e.g. aetiology (pregnancy, transfusion of blood products and/or previous transplant); number and time point], in disease context, and in immunosuppression regimens within the examined patient cohort. We overcame these limitations by studying HLA-specific alloantibody development in a unique patient cohort comprising healthy females that received a single intradermal injection of peripheral blood lymphocytes obtained from their partner, as part of their treatment for infertility with lymphocyte immunotherapy (LIT). The cohort comprised 191 couples with a median (SD) age of 34 (3) for females and 37 (4) for males. Comparison of male (henceforth referred to as donor) and female (henceforth referred to as recipient) HLA types revealed that the patient cohort was highly mismatched with a median number of 8 (IQR: 6-9) out of possible 12 HLA class I and class II mismatches (Figure 2). HLA mismatches between individual donor-recipient pairs were pooled and analysed together, accounting for potential effects at the patient level (see methods). After exclusion of HLA alleles that could not be determined at two-field level (donor HLA alleles n=38; recipient HLA alleles n=22), HLA mismatches where preformed donor specific antibody was identified on pre-LIT antibody screening with Luminex solid-phase assays (n=9), and HLA mismatches that were not represented in the Luminex single-antigen-bead panel (n=176), 1381 HLA mismatches were considered for further analyses (242 HLA-A, 266 HLA-B, 213 HLA-C, 257 HLA-DRB1, 247 HLA-DQ, and 156 HLA-DP).

**Figure 2.**
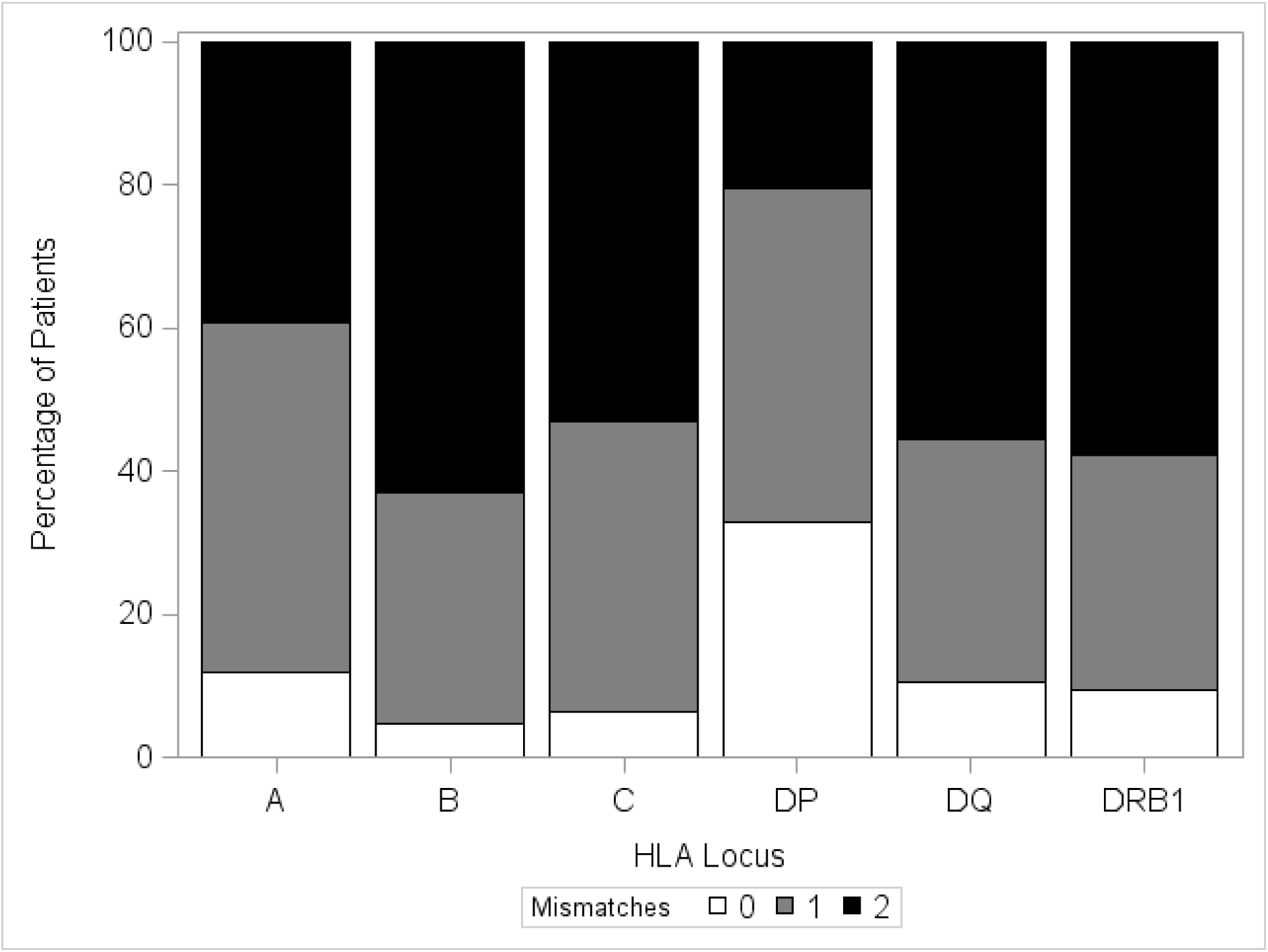
Distribution of HLA mismatches in the lymphocyte immunotherapy patient cohort. This figure shows the percentage of patients (lymphocyte donors) with 0, 1 or 2 mismatches within HLA-A, -B, -DRB1, -DQ and -DP loci.

Following LIT, HLA-specific antibody detection using Luminex showed that antibody binding against mismatched donor HLA had a median (IQR) MFI of 1039 (42, 5548). The majority of immunised recipients (84%) developed an IgG DSA response [defined as Mean Fluorescence Intensity (MFI) ≥2000] against one or more mismatched HLA expressed on their partner’s lymphocytes. Overall, DSA was detected against 569 of the 1381 (41%) donor-recipient HLA mismatches with a median (IQR) MFI of 6939 (3795, 9917). Luminex detected binding against donor HLA-C and -DP mismatches was of low magnitude [median (SD) MFI of 40.0 (1115.0) for HLA-C and 102.4 (2868.2) for HLA-DP] and DSA responses were less frequent [DSA against 12 of 213 (6%) donor HLA-C mismatches and against 32 of 156 (21%) donor HLA-DP mismatches] compared to other loci (Figure 3). Strong DSA responses (in frequency and magnitude) were noted against mismatches at HLA-A [160 of 242, 66%; median (IQR) MFI: 7610 (5406, 10295)], -B [138 of 266, 52%; median (IQR) MFI: 6852 (3546, 9353)], and -DQ [136 of 247, 55%; median (IQR) MFI: 7853 (4576, 11760)] loci followed by development of DSA against donor HLA-DRB1 [91 of 257, 35%; median (IQR) MFI: 4493 (3466, 9264)] alloantigens (Figure 3).

**Figure 3.**
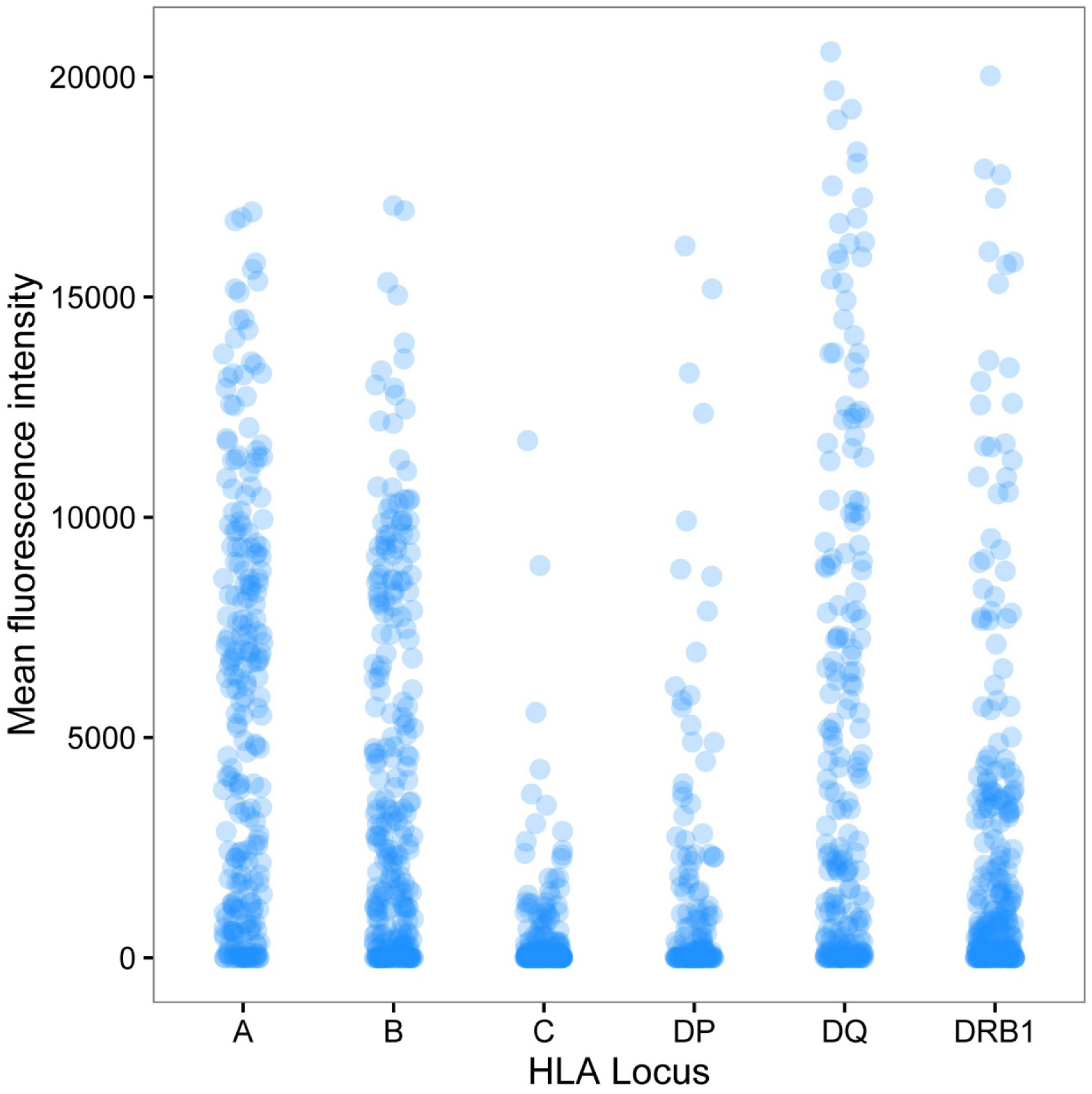
Donor-specific alloantibody responses after lymphocyte immunotherapy against mismatched HLA expressed on donor lymphocytes. The figure depicts alloantibody binding, as detected using Luminex single HLA beads, against mismatched HLA expressed on donor lymphocytes for the entire cohort. Donor-recipient HLA mismatches (n=1381) are grouped according to HLA locus (242 HLA-A, 266 HLA-B, 213 HLA-C, 156 HLA-DP, 247 HLA-DQ, and 257 HLA-DRB1) and the mean fluorescence intensity (MFI) of donor-specific antibody binding detected in recipient sera is shown on the y-axis. The median (SD) MFI of antibody responses against mismatches within individual HLA loci was 5270.3 (4625.6) for HLA-A; 2303.8 (4091.9) for HLA-B; 40.0 (1115.0) for HLA-C; 102.4 (2868.2) for HLA-DP; 2372.4 (5435.0) for HLA-DQ; and 803.1 (4056.2) for HLA-DRB1.

We examined the association between development of DSA and the immunogenicity of donor HLA mismatches as determined by comparative assessment of electrostatic potential between donor and recipient HLA (EMS-3D). Figure 4 shows the frequency of donor-recipient HLA mismatches according to their EMS-3D grouped by HLA locus; overall, the median (IQR) EMS-3D was 0.30 (0.24-0.35) and 0.22 (0.18-0.32) for HLA class I and class II, respectively. DSA responses against HLA-C mismatches were infrequent (supplementary Figure S4), reflecting the relatively low expression of HLA-C on lymphocytes (31), and were therefore not further evaluated. Logistic regression analysis showed that increasing EMS-3D of donor HLA was strongly associated with higher risk of DSA development (OR: 1.70 per 0.1 unit increase, 95% CI: 1.35-2.15, p<0.0001 for HLA-A and -B; OR: 2.56 per 0.1 unit increase, 95% CI: 2.13, 3.07, p<0.0001 for HLA-DRB1, -DQ, and -DP; Table 1). HLA-DP was the least immunogenic locus (excluding HLA-C) and it was notable that differences in electrostatic potential among HLA-DP mismatches were of lower magnitude compared to other loci (EMS-3D median: 0.19, IQR: 0.18-0.24), although this finding might in part reflect a lower cell surface expression of HLA-DP on lymphocytes compared to -DR and -DQ molecules (32-35). In contrast, physicochemical differences between donor and recipient HLA-DQ were higher compared to other loci (EMS-3D median: 0.35, IQR: 0.20-0.42) and donor HLA-DQ with the highest EMS-3D (within the fourth compared to first quartile), were highly likely to induce a DSA response (OR: 27.7, 95% CI: 10.4 - 73.9, p<0.0001). HLA-DRB1 mismatches had lower EMS-3D scores (EMS-3D median: 0.20, IQR: 0.17-0.24) and were less immunogenic compared to HLA-DQ, similar to what has been reported on the relative frequency of HLA-DRB1 and -DQ DSA responses after renal transplantation (14).

**Table 1.** Influence of donor HLA immunogenicity, as assessed by EMS-3D, and risk of development of donor-specific alloantibody after lymphocyte immunotherapy. EMS-3D: Electrostatic Mismatch Score - three dimensional; DSA: Donor-specific alloantibody; MM: mismatches Logistic regression models were used to investigate the association between donor HLA EMS-3D and donor-specific alloantibody development. Odds ratios are omitted where natural cubic splines were fitted.

| *HLA Locus* | **Odds ratio on developing HLA donor-specific antibodies (MFI≥2000)** |  |  |
| --- | --- | --- | --- |
|  | **Donor HLA MM, DSA** |  |  |
| ***HLA-A*** | **N=242, 160 events** | **OR (95% CI)** | **p-value** |
| **EMS-3D** |  | 1.64 (1.14, 2.37) | 0.007 |
| ***HLA-B*** | **N=266, 138 events** | **OR (95% CI)** | **p-value** |
| **EMS-3D** |  | 1.57 (1.15, 2.15) | 0.004 |
| ***HLA-DRB1*** | **N=257, 91 events** | **OR (95% CI)** | **p-value** |
| **EMS-3D** |  |  | <0.0001 |
| ***HLA-DQ*** | **N=247, 136 events** | **OR (95% CI)** | **p-value** |
| **EMS-3D** |  | 2.64 (2.01, 3.48) | <0.0001 |
| ***HLA-DP*** | **N=156, 32 events** | **OR (95% CI)** | **p-value** |
| **EMS-3D** |  | 3.32 (1.41, 7.81) | 0.004 |
| ***HLA-A+B*** | **N=508, 298 events** | **OR (95% CI)** | **p-value** |
| **EMS3D** |  | 1.70 (1.35, 2.15) | <0.0001 |
| ***HLA-DP+DQ+DRB1*** | **N=660, 259 events** | **OR (95% CI)** | **p-value** |
| **EMS3D** |  | 2.56 (2.13, 3.07) | <0.0001 |
| ***HLA-A+B+DP+DQ+DRB1*** | **N=1168, 557 events** | **OR (95% CI)** | **p-value** |
| **EMS3D** |  | 2.35 (2.04, 2.71) | <0.0001 |

**Figure 4.**
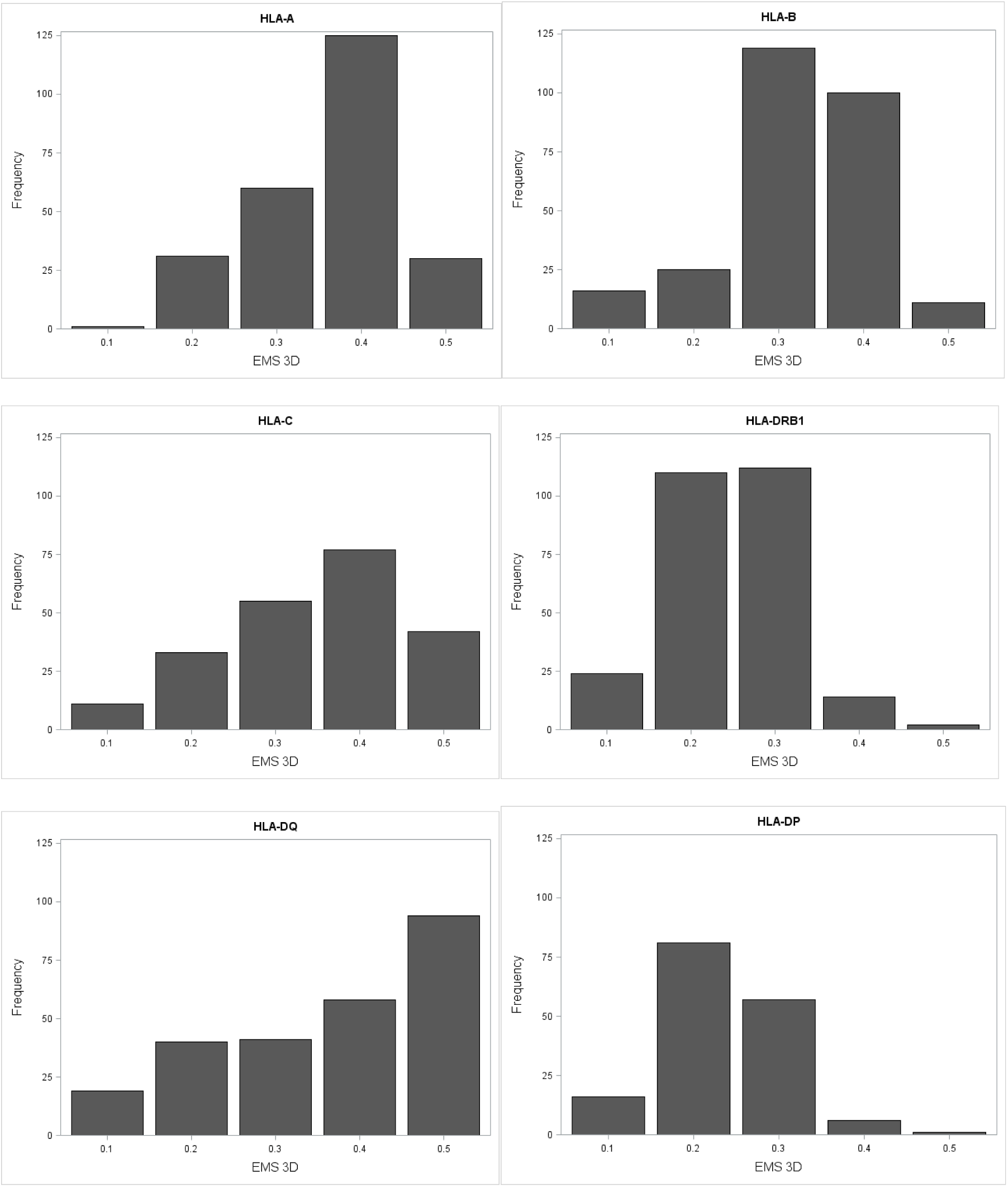
Frequency of HLA class I and class II mismatches in the lymphocyte immunotherapy patient cohort according to their Electrostatic Mismatch Score (EMS-3D). The figure depicts the frequency of donor-recipient HLA mismatches according to their EMS-3D grouped by HLA locus. The median (IQR) EMS-3D for individual loci was HLA-A: 0.32 (0.27-0.36); HLA-B: 0.28 (0.22-0.33); HLA-C: 0.32 (0.22-0.40); HLA-DRB1: 0.20 (0.17-0.24); HLA-DQ: 0.35 (0.20-0.42); and HLA-DP: 0.19 (0.18-0.24).

The potential of this approach to predict the immunogenic potential of donor HLA was examined further by considering the probability of an alloantibody response after LIT according to the EMS-3D of donor HLA alloantigens, using logistic regression modelling. As shown in Figure 5, this analysis demonstrated a strong relationship between donor-recipient HLA electrostatic disparity and predicted probability of an alloantibody response for all loci examined. Wider confidence intervals were observed for the immunogenic potential of high EMS-3D HLA-DR and -DP alloantigens and this reflected the relatively low number of observations for such mismatches in the patient cohort. The association was strongest for HLA-DQ alloantigens (Figure 5D) and it is notable that multiple recent studies have highlighted the predominance of HLA-DQ specific humoral alloresponses after solid organ transplantation (36-39). Overall, our model predicts that donor HLA class I and class II (HLA-A, -B, -DRB1, -DQ, and -DP) with low EMS-3D have an approximately 10% probability of inducing DSA [e.g. the observed probability of an alloantibody response for HLA with EMS-3D<0.045 was 11% (of 27 mismatched HLA 3 induced DSA)]. This probability increases to over 75% for donor HLA with the highest EMS-3D [e.g. the observed probability of a DSA response for donor HLA with EMS-3D>0.38 was 71% (of 110 mismatched HLA 78 induced DSA)]. Fitting the model to examine high level DSA responses (defined as DSA MFI≥8,000 which, in the context of transplantation, commonly denotes high immunological risk) showed a near linear association between donor HLA EMS-3D and predicted probability of DSA development (Figure 5G).

**Figure 5.**
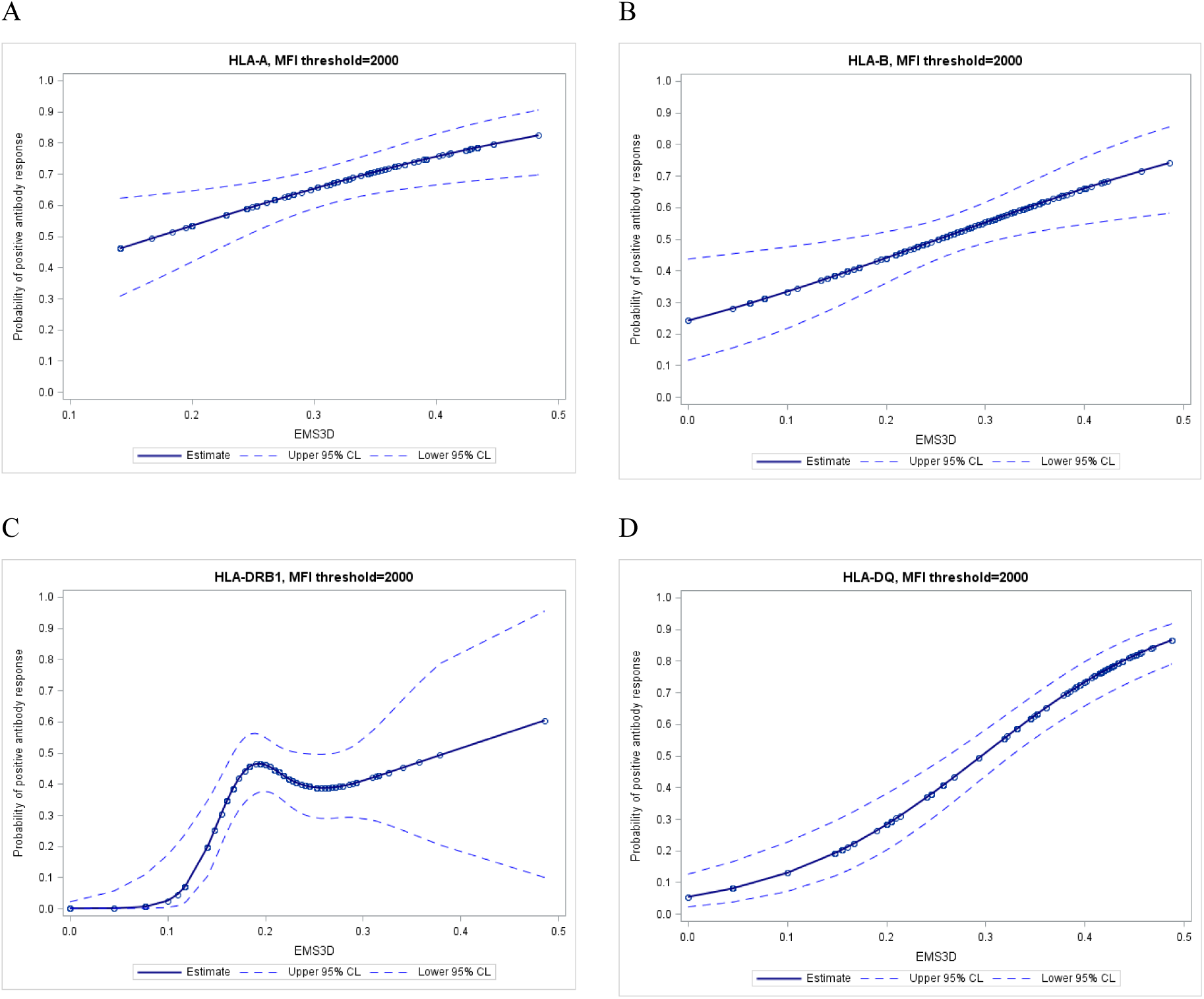
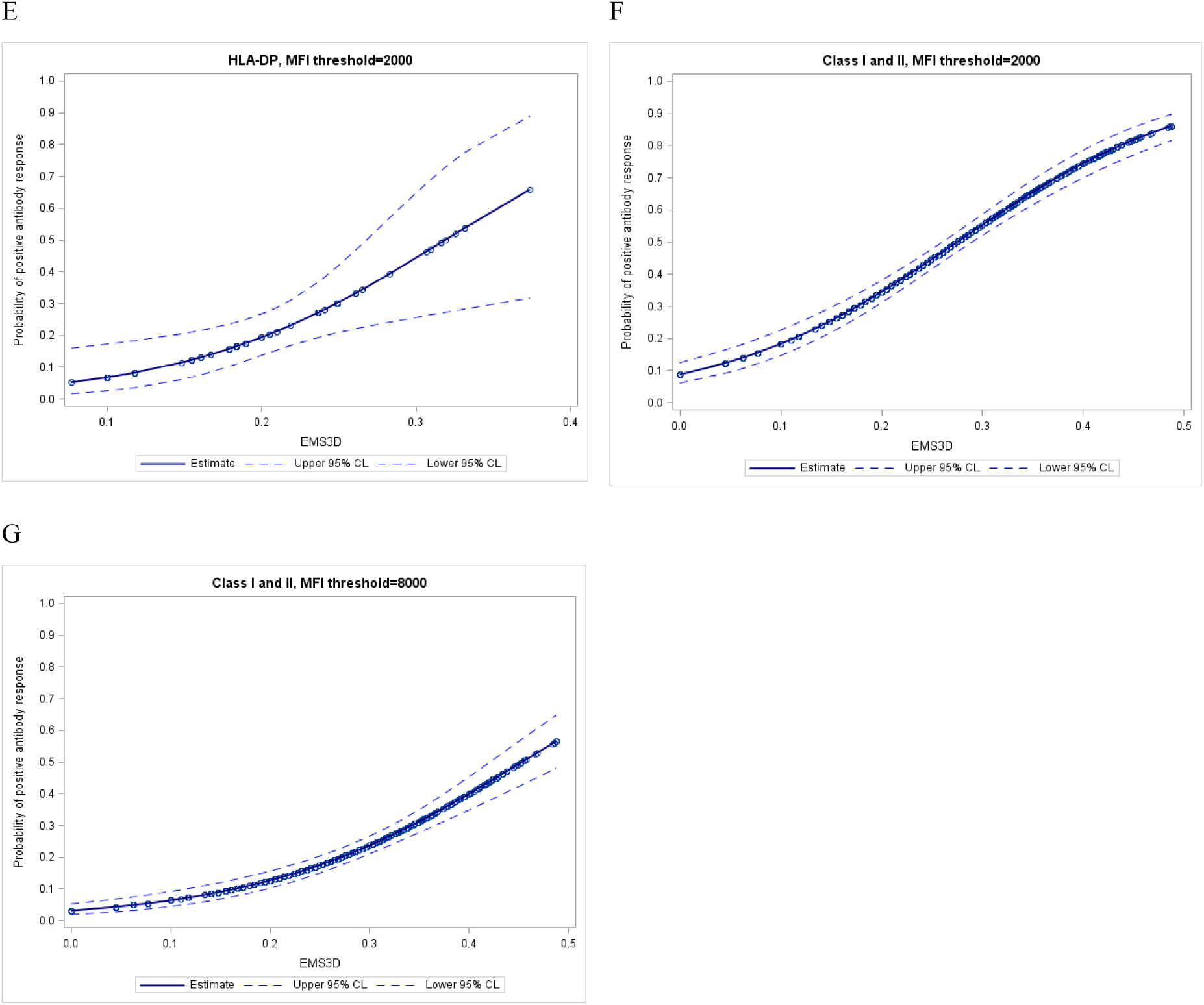
Probability of donor-specific alloantibody response after lymphocyte immunotherapy according to the Electrostatic Mismatch Score (EMS-3D) of mismatched HLA on donor lymphocytes. The relationship between the immunogenic potential of donor HLA, as determined by EMS-3D, and the probability of a donor-specific alloantibody response after lymphocyte immunotherapy was examined using logistic regression modelling. Each panel shows a logistic regression model with 95% confidence intervals (dotted lines) for individual HLA loci (panels A-E) and for HLA class I and class II loci combined (panels F and G). DSA responses against HLA-C mismatches were infrequent and were not examined. Donor-specific alloantibody responses were defined using mean fluorescence intensity (MFI) cut-off thresholds of ≥2,000 (panels A-F) and of ≥8,000 (panel G). Wide confidence intervals for alloantibody responses against HLA-DR and -DP alloantigens reflect the relatively low number of observations for HLA-DR and -DP mismatches with high EMS-3D scores in the lymphocyte immunotherapy patient cohort. Relatively few alloantibody responses with MFI≥8,000 were noted against HLA-DP alloantigens (n=7) and, therefore, HLA-DP mismatches were not included in the panel G model.

We next considered the relationship between donor HLA EMS-3D and the magnitude of the alloantibody response as assessed based on the MFI binding detected in the Luminex assay. The latter provides semi-quantitative information on the level of circulating alloantibody and previous studies have shown an association between DSA MFI level and clinical outcome (5, 40, 41). Median regression analysis showed that donor HLA with increasing EMS-3D were associated with progressively stronger (higher MFI) alloantibody responses following LIT (p<0.001; Figure 6). The magnitude of the alloantibody response increased from a median MFI of 48 (IQR: 0-300) for donor HLA class I and II (HLA-A, -B, -DRB1, -DQ) with EMS-3D <0.14 to a median MFI of 6432 (IQR: 1876-10002) for alloantigens with EMS-3D>0.35 (Figure 6). The association between donor HLA EMS-3D and MFI binding level was strongest for donor HLA class II mismatches (Figure 6B).

**Figure 6.**
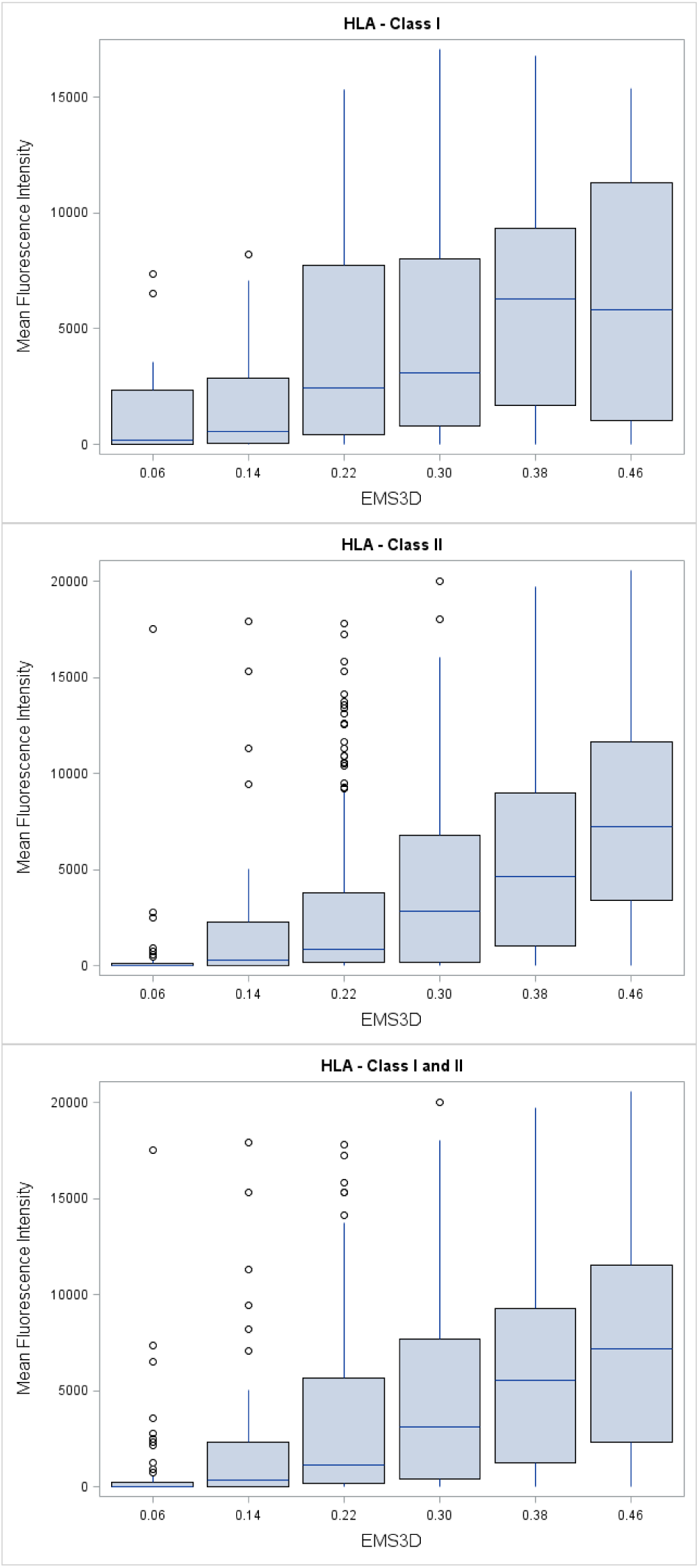
Relationship between donor HLA Electrostatic Mismatch Score (EMS-3D) and the donor-specific Mean Fluorescence Intensity (MFI) binding level of alloantibodies developed after lymphocyte immunotherapy. The relationship between EMS-3D of mismatched HLA on donor lymphocytes and the magnitude of donor-specific antibody binding, as assessed based on mean fluorescence intensity (MFI) detected in the Luminex single antigen bead assay, is depicted. Donor HLA mismatches are grouped according to EMS-3D and the box plots depict the median MFI (horizontal blue line) and interquartile range (box) of MFI values (the lines show maximum MFI values) for donor-specific antibody binding within each group. Median regression analysis showed that donor HLA with increasing EMS-3D were associated with progressively stronger (higher MFI) alloantibody responses following lymphocyte immunotherapy (p<0.001). Alloantibody responses against donor HLA-C and -DP mismatches were infrequent and of low MFI value and are, therefore, not included in this analysis.

Donor homozygocity for a given HLA class I or II mismatch had no effect on the risk of DSA development (data not shown). Although there was evidence that alloantibody responses were more likely the higher the amount of donor lymphocytes administered during LIT (adjusted OR: 1.02 per 10^6^ increase in lymphocyte dose, 95% CI: 1.01-1.03, p<0.0001), adjusting for donor lymphocyte dose did not alter the relationship between donor HLA EMS-3D and risk of DSA development. In the analyses above, donor-recipient HLA mismatches and their immunogenic potential were considered individually and independently from each other. To consider HLA immunogenicity at a locus and individual patient level, we examined the association between the overall immunogenic potential of HLA mismatches within a locus (as assessed by EMS-3D) and the likelihood of a recipient locus-specific DSA response (as suggested by other authors (14, 42)). This analysis showed that locus-specific EMS-3D was strongly associated with HLA-A, -B, -DRB1, -DQ, and -DP DSA development (OR: 1.40 per 0.1 unit increase, 95% CI: 1.21-1.61, p<0.0.001 for HLA-A and -B; OR: 1.57 per 0.1 unit increase, 95% CI: 1.40-1.77, p<0.0001 for HLA-DR, -DQ, and -DP) independent of lymphocyte dose administered during LIT.

### Analysis of HLA-specific antibody responses after kidney transplantation

We next considered, in a proof of principle study, the applicability of our approach in the kidney transplantation setting. Alloantibody responses against donor HLA expressed on renal allografts were examined in a cohort of 131 kidney transplant recipients returning to the transplant waiting list following first graft failure. Humoral responses against HLA class II alloantigens predominate long term after kidney transplantation and are strongly associated with graft failure (1) and, therefore, this analysis focused on development of DSA against mismatched HLA class II alloantigens. The demographic and transplant characteristics of the patient cohort have been published previously (6). To account for factors that may influence HLA-specific antibody responses at an individual patient level, multivariable logistic regression analysis of the association between EMS-3D of donor kidney HLA-DRB1 and -DQ mismatches and DSA development was adjusted for length of time to graft failure, length of time on the waiting list after listing for re-transplantation, maintenance immunosuppression regimen whilst on the transplant waiting list, graft nephrectomy and number of administered blood transfusions. Similar to the findings on DSA responses after LIT, this analysis showed that the immunogenicity of HLA-DRB1 and -DQ mismatches expressed on donor kidneys, as assessed by their EMS-3D, was an independent predictor of development of DSA (OR: 1.90 per 0.1 unit increase, 95% CI: 1.25-2.88, p=0.0026 for HLA-DRB1; OR: 1.86 per 0.1 unit increase, 95% CI: 1.07-3.23, p=0.028 for HLA-DQ).

## Discussion

The capacity of donor HLA to stimulate alloantibody responses (HLA immunogenicity) is dependent upon their structural recognition by receptors on recipient B cells that initiate the immune response, and previous work has suggested that HLA immunogenicity should be considered in the context of amino acid sequence polymorphisms between donor and recipient HLA molecules (12, 21, 43). The present investigation introduces a fundamentally new approach at evaluating the immunogenic potential of donor HLA focusing entirely on their tertiary structure and on their unique structural and surface electrostatic potential properties compared to recipient HLA molecules. We have developed a computational approach to compare and quantify HLA electrostatic properties at atomic resolution level and applied it to predict HLA-specific alloantibody development in a unique model of human sensitisation. We show that (a) HLA molecules differ widely at the level of electrostatic potential in three-dimensional space and these differences are not explicable on account of the underlying amino acid sequence polymorphisms; (b) the electrostatic disparity of a donor HLA compared to recipient HLA molecules, as assessed by EMS-3D, was strongly associated with the risk of development of donor-specific alloantibody; and (c) electrostatic potential disparities are highest among HLA-DQ molecules which were the most immunogenic alloantigens in this study and whose immunogenicity conformed best to our EMS-3D algorithm. Taken together with our proof of principle study in the setting of human kidney transplantation, the present investigation provides important first validation that donor HLA immunogenicity can be predicted based on assessment of their unique surface electrostatic potential properties compared to recipient HLA molecules.

The risk of allosensitisation after transplantation increases incrementally with the number of HLA mismatches at individual HLA class I and class II loci (6). However, simple enumeration of differences at the whole antigen level is constrained by limited possible values (zero, one or two mismatches per locus) and does not account for differences in donor HLA immunogenicity for a given recipient.

Current approaches for determining the potential of a donor HLA to induce an alloantibody response are based on quantifying the degree of ‘dissimilarity’ between the donor and recipient HLA molecules (12, 17, 19, 21). The most frequently used methods (HLAMatchmaker and Cambridge HLA immunogenicity algorithm) evaluate differences in the number and location of amino acid mismatches at continuous and discontinuous (eplets) positions on the HLA sequence and multiple studies have suggested they provide superior risk stratification over conventional HLA mismatch grade for predicting development of DSA, allograft rejection, transplant glomerulopathy and allograft survival (14, 16, 18, 20, 42). Both of these methods reflect differences in amino acid sequence between donor and recipient HLA mismatches and generate highly correlated scores that provide a similar assessment of HLA immunogenicity (18). Importantly, accounting for the physicochemical properties of donor HLA amino acid polymorphisms appears to improve prediction of DSA development against HLA class I alloantigens (18, 19).

Protein electrostatics reflect the amino acid composition of their primary structure but are mainly determined by the number and distribution of polar and charged residues, the protonation state of ionisable groups within a given ionic environment, and their ability to form specific bonding interactions such as salt bridges and hydrogen bonds. Importantly, our study shows that the variation in surface electrostatic potential between HLA molecules cannot be inferred on account of differences at the amino sequence level and relatively poor correlation exists between residue polymorphisms and electrostatic disparity among HLA class I and class II molecules (supplementary Figure S2). Given that hydrophobic patches on a protein surface tend to have low electrostatic potential, compared to an acidic, basic or polar patch, the electrostatic potential also captures aspects of the protein’s hydrophobic interaction properties. Electrostatic forces are important determinants of the affinity and specificity of macromolecular interactions and it has been suggested that the process of affinity maturation involves optimisation of electrostatic interactions in the B cell receptor-antigen binding site (24, 25, 44, 45). Our study showed that donor HLA with high versus low EMS-3D were more likely to induce a specific alloantibody response and, therefore, it would be tempting to speculate that alloantigens with disparate electrostatic potential profiles compared to recipient HLA molecules lead to more efficient B cell receptor recognition in the secondary lymphoid organs and to improved selection and survival of differentiated B cells during the process of affinity maturation in the germinal centre. Indeed, recent insights into the mechanisms that determine the fate decision of proliferating, antigen-activated B cells at the pre-germinal centre stage suggested that B cells with higher affinity to their antigen presented more HLA-peptide to and made longer-lasting contact with cognate T follicular helper (T_FH_) cells at the B cell - T cell border in secondary lymphoid organs, resulting in more T cell help and differentiation into germinal centre B cells (with further B cell receptor diversification through somatic hypermutation) (46, 47). In contrast, proliferating B cells with lower affinity to their antigen may form less durable T_FH_ cell - B cell conjugates and are more likely to develop into germinal centre-independent memory B cells (that undergo class switching but not somatic hypermutation) (48). The implication of this model of antigen-activated B cell differentiation for the present investigation is that alloantibody responses to donor HLA with high EMS-3D might be derived by germinal centre-dependent B cells and are of high affinity whereas humoral responses to donor HLA with lower EMS-3D, when triggered, might be derived by germinal centre-independent B cells and are more broadly reactive and of lower affinity. It would be interesting to investigate this hypothesis in future studies.

Multiple studies over recent years have provided strong evidence in support of the association between the development of donor HLA-specific alloantibodies and the risk of acute antibody-mediated rejection, chronic rejection and allograft loss across all solid organ transplants (42, 49-54). The ability to assess the risk of post-transplant humoral alloimmunity associated with particular donor-recipient HLA combinations is of major clinical interest both to inform organ allocation policies and to enable more efficient immune monitoring and individualisation of immunosuppression protocols to help prevent *de novo* DSA development. The present study suggests that the probability of an alloantibody response (generation and magnitude) against a donor HLA-A, -B, -DRB1, -DQ or -DP alloantigen increases with increasing EMS-3D and that it is possible to identify a substantial number of low EMS-3D HLA mismatches that might be tolerated by the immune system of a given recipient. DSA development against both HLA class I and class II alloantigens increases the risk of subsequent rejection and allograft failure, but humoral responses against HLA class II seem to predominate and these most commonly involve HLA-DQ specific alloantibodies (37-39, 55). Our analysis of the most common HLA-DQ alleles (supplementary Figure S2) showed that electrostatic potential disparities are highest among HLA-DQ alloantigens (with only a modest correlation between ESD and the underlying amino acid sequence polymorphism) compared to other loci and this may account for their increased immunogenic potential. Notably, HLA-DQ immunogenicity conformed best to our prediction model with a strong association between donor EMS-3D and probability of a DSA response in the LIT cohort, whereas a strong association between HLA-DQ EMS-3D and DSA development was also noted in the kidney transplant cohort.

The present study has focused on the structural aspects of donor HLA allorecognition that influence the subsequent humoral response by recipient B cells. Our computational protocol, enables quantification of electrostatic potential differences between donor and recipient HLA accounting for the entire three-dimensional space around the HLA molecule to produce an average score. Exclusion of the membrane bound area of the HLA did not alter the results of our analysis as this part of the molecule is relatively monomorphic and therefore similar among different HLA. However, it is possible that a small surface area on a donor HLA that differs widely in electrostatic potential from the respective area on recipient HLA might be sufficient to drive the alloimmune response, even though the average difference across the entire molecule remains low. Our computational method can be adapted to incorporate immunogenic ‘hot-spots’ on the HLA molecular surface (e.g. functional B cell epitopes) and this is the subject of our current research (28). It is also important to recognise that proliferation and differentiation of antigen-specific naïve B cells into memory B cells and long-lived plasma cells requires T cell help through linked recognition of antigen derived peptides presented in the context of B cell HLA class II molecules. Previous observational studies highlighted the importance of the HLA-DR phenotype of the recipient in humoral alloresponses to donor HLA class I alloantigens (56, 57) and, more recently, this concept has been extended to evaluate the capacity of recipient HLA-DR molecules to bind donor HLA class I and class II derived peptides using the netMHCIIpan computational method and to examine the contribution of this pathway to DSA development (58-61). Computational prediction of HLA class II restricted epitopes by CD4^+^ T cells is of great interest for understanding immune responses in the context of transplantation, autoimmunity, infection and cancer but it is a difficult and complex undertaking due to the open conformation of the HLA class II peptide binding groove that can accommodate peptides of variable length (10 to 30 amino acids long) and due to the fact that antigen processing and peptide loading are incompletely understood (62). Nevertheless, this is the subject of intense research and a multitude of computational methods are currently available for predicting HLA class II peptide binding, with mixed results (63, 64). Despite the strong association between the surface electrostatic potential properties of donor HLA and alloantibody responses in the current study, DSA development was noted against HLA with low EMD-3D and *vice versa*. Consideration of HLA class II peptide presentation to CD4^+^ T cells is likely to improve the predictive ability of our immunogenicity algorithm and we are currently undertaking relevant studies to explore this question. To this extent, it would be intriguing to examine the contribution of electrostatic interactions within the pockets of the HLA class II peptide binding groove to high affinity peptide binding (65, 66).

A strength of the present study is that HLA immunogenicity was investigated in a unique model of HLA sensitisation comprising non-sensitised individuals that underwent a single sensitising event (injection of donor lymphocytes) followed by detection of HLA-specific development approximately five weeks later. To our knowledge, this is the first study to systematically examine HLA immunogenicity in this setting, thus avoiding commonly encountered confounders in similar studies including variations in baseline disease, non-uniform immunosuppression protocols, and unknown and variable sensitisation events (previous transplants, pregnancies and blood transfusions). We acknowledge that it would have been interesting to examine the temporal evolution of the immune response at different time points but this was not possible as further serum samples were not routinely collected. The kinetics of IgG HLA-specific antibody development after exposure to human lymphocytes have not been studied in detail, especially using sensitive detection methods, but it is well documented that specific humoral responses (class switched and affinity matured) peak within 30 days from exposure to antigen (48), and a previous study in the context of LIT suggested that the alloresponse peaks during the second month post lymphocyte immunisation (67). It is, therefore, unlikely that relevant alloantibody responses have been missed due to the timing of serum collection in this study. Another limitation of the study is that a relatively small cohort of patients was examined to investigate development of DSA after failure of kidney transplant and our findings would be strengthened if they were confirmed in larger cohorts and in patients with functioning grafts that are prospectively monitored for alloantibody development, accounting for immunosuppression regimen, drug levels and for non-compliance. We were unable to systematically assess the immunogenicity of HLA-C and -DP alloantigens due to their relatively low expression on lymphocytes. Alloantibodies against HLA-DP can cause humoral rejection after kidney transplantation and mismatching at the -DP locus is associated with decreased graft survival in patients undergoing repeat kidney transplantation (68, 69). Within the confines of the lymphocyte HLA sensitisation model, HLA-DP immunogenicity seemed to conform to our algorithm and it would be interesting to examine the applicability of our approach in the transplant setting both in terms of predicting HLA-DP specific sensitisation and for analysis of the relevant effect of -DP mismatching on renal transplant outcomes.

In conclusion, the present investigation demonstrates a clear relationship between the electrostatic properties of HLA molecules and their immunogenic potential. Quantification of electrostatic potential differences at the tertiary level between donor and recipient HLA molecules enables prediction of humoral alloimmune responses in the context of lymphocyte allosensitisation. We have shown the translational potential of this approach in the clinical setting of kidney transplantation. Our approach has the potential to identify acceptable HLA mismatches for a given recipient, thereby increasing access to suitable donors that are currently considered a poor match, to enable better immunological risk assessment, and to decrease the burden of allosensitisation and humoral rejection after solid organ transplantation.

## Methods

### Lymphocyte immunotherapy patient cohort

The study cohort comprised women who had been referred to the Institute of Immunology, University Schleswig-Holstein, Campus Kiel, Germany for lymphocyte immunotherapy (LIT) as treatment for infertility in patients with recurrent first trimester miscarriage and/or patients with recurrent embryo implantation failure after *in vitro* fertilisation. LIT was introduced at the Institute of Immunology, University Schleswig-Holstein, Campus Kiel, Germany in the 1980s, is approved by the local health authority and is covered by the national health insurance providers (70, 71). LIT comprised a single intradermal injection of lymphocytes isolated from 50mls of peripheral venous blood obtained from their male partner. The lymphocytes were separated under sterile conditions by Ficoll-Hypaque density gradient centrifugation and after two washing steps the cells were re-suspended in 1 ml of normal saline for injection. The cellular content of the suspension was examined using phase contrast microscopy (average cellular contents have been reported previously (71)). The suspension, without prior storage, was given to the female partner by intradermal injection at the volar side of one forearm (71, 72). Peripheral blood was collected from all women within 2 months before LIT and 5 weeks (median: 33 days, SD: 4.5) following LIT and serum stored at −21°C for subsequent detection of antibodies to HLA. DNA was also isolated from peripheral blood of all women and their partners for HLA typing.

This retrospective study comprised 191 consecutive women (and their male partner) who underwent their first LIT in 2009 and 2010 and had not had a previous pregnancy, blood transfusion, or organ transplantation, were not on immunosuppressive medication and had no detectable antibodies directed against their partners’ HLA, as defined by complement-dependent cytotoxicity assays. During LIT the cohort received a median (SD) of 37.4 (15.0) x 10^6^ of their partner´s lymphocytes (range 20.2 - 83 × 10^6^ lymphocytes).

### HLA typing

DNA samples were genotyped using the Immunochip®, an Illumina iSelect HD custom genotyping array according to Illumina protocols at the Institute of Molecular Biology, Kiel University. Genotype calling was performed using Illumina´s GenomeStudio Data Analysis software and the custom-generated cluster file of Trynka et al (73) based on an initial clustering of 2,000 UK samples with the GenTrain2.0 algorithm and subsequent manual readjustment and quality control. Subsequent imputation of classical HLA alleles from SNP genotypes was performed using two independent HLA imputation pipelines, HLA*IMP2 (74) and SNP2HLA (75). HLA-A and -B typing was also performed (as part of the LIT protocol) using a reverse polymerase chain reaction sequence specific oligonucleotide system as implemented in the Luminex platform (LABType® SSO, One Lambda, Canoga Park, CA, USA) and the results used for quality control in case of ambiguous results. Missing genotype data from failed genotype calls or failed quality control (n=191) were imputed, where possible, using an allele frequency-based prediction tool that considers HLA haplotype and patient race (13) (n=39 HLA class I alleles and n=92 HLA class II alleles) or excluded from further analysis (n=60 HLA class II alleles).

### HLA-specific antibody screening

Serum samples obtained before and after LIT were screened for HLA-specific antibodies using solid-phase Luminex HLA antibody-detection beads (LABScreen, One Lambda, Canoga Park, CA). Selected HLA-specific antibody-positive samples were analysed using Luminex single-antigen HLA class I and class II antibody-detection beads (One Lambda, Canoga Park, CA). HLA single-antigen bead defined antibody reactivity was determined using mean fluorescence intensity (MFI) cut-off thresholds of 2,000 (MFI cut off level used clinically in our Centre and elsewhere to define a positive alloantibody response to a given HLA), to denote the presence of donor-specific antibodies (DSA), and of 8,000 to reflect high DSA levels (widely accepted Luminex MFI level at which DSA often results in a positive donor complement dependent cytotoxicity crossmatch test; DSA above this level commonly denotes higher immunological risk in the context of transplantation).

### Comparative structure modelling of HLA alleles

Comparative structure models of all HLA class I and class II alleles represented in the HLA types of the patient cohort and of all common HLA alleles (frequency >1%) were generated using the program MODELLER v9.17 (https://salilab.org/modeller/) (76). Templates for comparative structure modelling were identified by querying the RCSB Protein Database using the sequence of HLA-B*07:02 and HLA-DRB1*01:01, for HLA Class I and HLA Class II, respectively. The search was carried out using the DELTA-BLAST algorithm, for humans (Taxonomy ID: 9606, E-value threshold of 0.005) and identified 125 HLA Class I and 41 HLA Class II unique crystallographically resolved structures. Of these, 12 HLA class I structures (PDB codes: 1K5N, 3MRE, 3CZF, 3BWA, 3LN4, 3SPV, 2BVP, 2A83, 3MRB, 1X7Q, 1OGT, 1XH3) and 22 HLA class II structures (PDB codes: 4P4R, 4P57, 4P5K, 4P5M, 1JK8, 1UVQ, 1S9V, 2NNA, 4MD4, 4MD5, 4MDJ, 3PDO, 3C5J, 1FV1, 1AQD, 1T5W, 2Q6W, 3L6F, 3QXD, 4H25, 4I5B, 4OV5) were retained for comparative modelling based on favourable indices of structural quality (Ramachandran plot, R factor, crystallographic resolution, DOPE, Verfiy3D, PROCHECK and WHAT_CHECK scores) (77-80) and following exclusion of HLA structures resolved in complex with a ligand, such as T-cell or KIR-receptors (to avoid potential conformational distortion of the HLA structure occurring upon binding to the ligand). The sequences of the extracellular domain of target HLA molecules were retrieved from the EBI sequence database (ftp://ftp.ebi.ac.uk/pub/databases/ipd/imgt/hla/) and aligned using Clustal W2 using the *BLOSUM* matrix and the *Neighbor Joining* clustering algorithm and manually adjusted as indicated (81). Mean sequence homology between templates and target sequences was 91.9% (range: 84.1 - 100.0%). To standardise the peptide binding groove environment and eliminate structural variations between modelled molecules due to the peptide sequence, all HLA class I and class II structures were modelled with an alanine nonamer peptide and an alanine 12-mer peptide, respectively. The integrity of the modelled structures was validated using multiple objective measures of structural quality (Ramachandran plot, DOPE, Verfiy3D, PROCHECK and WHAT_CHECK scores; data not shown).

### Electrostatic potential calculations

The side chains of modelled HLA structures were protonated using PROPKA (82) and atom charges and radii were assigned using the PARSE force-field (83), as implemented in PDB2PQR (84), at physiological pH of 7.4. The electrostatic potential in three-dimensional (3D) space surrounding each HLA structure was calculated numerically by solving the linearised Poisson-Boltzmann equation using the finite difference/ finite element approach as implemented in APBS (http://www.poissonboltzmann.org/) for each point on a cubic grid with sides of 353 points at a spacing of 0.33 Å. Other parameters were set as follows: ionic solution of 0.15 M of univalent positive and negative ions; protein dielectric of 2; solvent dielectric of 78; temperature of 310 K; and a probe radius of 1.4 Å (28, 85).

### Quantitative comparison of 3-dimensional electrostatic potential between HLA molecules

Electrostatic potential comparisons between two HLA molecules of interest were performed based on the method described by Wade et al (http://pipsa.eml.org/pipsa/) (86, 87). As described previously (28, 66), the method considers the electrostatic potential in a region or ‘layer’ of space above the molecular surface of a protein. This ‘layer’ of thickness δ, is defined at a distance σ from the van der Waals surface of the protein and the electrostatic potential at the cubic grid points encompassed by this ‘layer’ are considered for subsequent calculations. Electrostatic potential comparisons between two HLA molecules of interest are performed for grid points within the intersection of their ‘layers’ after the two structures are superimposed (87), as shown in Figure 1C. Quantitative comparisons are performed using the Hodgkin similarity index (SI) (88) which assigns values between 1 (electrostatic identity, both in magnitude and sign) and −1 (electrostatic anti-correlation of the sign of the potential but of the same magnitude). These values are then converted into a distance [Electrostatic Similarity Distance 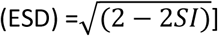 to give values between 0 (electrostatic identity) and 2 (electrostatic anti-correlation) where 1 represents no apparent correlation. For the purpose of this study, a ‘layer’ of δ=4 Å thickness and raised σ=3 Å above the molecular surface of HLA molecules was defined. These values were selected so that the electrostatic potentials compared were not highly sensitive to small changes in molecular structure (87). Electrostatic potential calculations considered the 3-dimensional space around the entire HLA molecule. Exclusion of the membrane-bound region of the HLA molecule from electrostatic potential comparisons had minimal impact on the scores (ESD) obtained (data not shown). Similarly, sampling the electrostatic potential in ‘layers’ of different δ thickness (1 Å - 5 Å) and different σ distance from the molecular surface (1 Å - 5 Å) did not alter the results of the quantitative comparisons (data not shown).

For HLA alleles within a locus, electrostatic potential comparisons were made in a pair-wise, all-versus-all fashion. The ESDs generated by the comparisons were compiled as a distance matrix that was then displayed as a symmetrical heat map and as a dendrogram with allele re-ordering such that electrostatically similar alleles cluster together. Symmetrical heat maps, dendrograms and allele re-ordering were performed in R using complete-linkage hierarchical clustering as implemented in the *hclust* function (89).

### Three-dimensional electrostatic mismatch score (EMS-3D)

The HLA type of the male partner (‘donor’) was compared to the HLA type of the female partner (‘recipient’) to identify mismatches in HLA-A, -B, -C, -DRB1, -DQ and-DP loci. As shown in supplementary Figure S1, for HLA class I mismatches, the donor HLA was electrostatically compared to each of the recipient HLA class I alloantigens to derive the respective ESDs and the minimum ESD value was taken to represent the 3-dimensional electrostatic mismatch score (based on the inter-locus comparison principle as previously described in our sequence based immunogenicity algorithm (21)). For HLA class II mismatches (supplementary Figure S1), the donor HLA was electrostatically compared to each of the recipient HLA within the same locus (intra-locus comparison) to derive the ESDs and the minimum value was taken to represent the EMS-3D (19).

### Transplant patient cohort

The patient population studied and the antibody screening protocol used have been described in detail previously (6). Briefly, the study cohort comprised 131 consecutive patients (87 males, 44 females, median age 38) who received a primary kidney allograft between 1995 and 2010, and returned to the Cambridge kidney transplant waiting list following failure of their graft during this time period [56 patients (43%) underwent transplant nephrectomy]. Approximately half (50.4%) of the patients in the cohort continued to receive immunosuppression after return to the waiting list (of those, 55% received single agent immunosuppression and 45% received dual agent immunosuppression). Antibody screening was undertaken at the time of (and prior to) the first transplant, after return to the transplant waiting list following graft failure and at 3 monthly intervals while remaining on the list for re-transplantation. Screening was undertaken using Luminex single antigen beads (One Lambda, Canoga Park, CA), as described above. The median (SD) duration of follow up since transplantation was 2539 (1605) days.

### Statistics

To investigate whether there was an association between donor HLA EMS-3D and DSA development, all HLA mismatches in the patient cohort were considered together using logistic regression models. Random effects were considered to account for potential correlations within individual patients, however these were not found to be important and not included in the final models. An MFI≥2000 was classified as a positive result for presence of DSA, and an MFI≥8000 as a high level DSA response. Where the donor was homozygous for a particular HLA mismatch, only one observation (mismatch) was included in the model. EMS-3D was modelled as a continuous variable and for illustration as a categorical variable by splitting into quartiles. To investigate HLA immunogenicity at an HLA locus and individual patient level, logistic regression models were used to examine the association between total EMS-3D for HLA mismatches within a locus (one or two mismatches) and development of a recipient locus-specific DSA response, adjusting for the effect of lymphocyte dose administered during LIT. Median regression models were used to model the effect of EMS-3D on post-LIT DSA MFI value. To assess non-linearity of the explanatory variable, EMS-3D, natural cubic spline terms were added to the logistic regression model and these terms were kept in the model if there was sufficient evidence of non-linearity. For analyses of the transplant cohort, logistic regression models were used to investigate the relationship between donor HLA EMS-3D and DSA development at the patient level, accounting for clinical explanatory variables. Initially, each explanatory variable was modelled separately; further models investigated the additional value in incorporating EMS-3D into models including length of time to graft failure, length of time on the waiting list after listing for re-transplantation, maintenance immunosuppression regimen, graft nephrectomy and number of administered blood transfusions. Models were compared using the log likelihood ratio statistic and p-values of 0.05 or less were considered significant. All analyses were conducted using SAS (version 9.4).

### Study approval

All participating couples in the lymphocyte immunotherapy cohort gave informed written consent prior to inclusion in the study for use of their data and blood samples for research and this study was approved by the local Institutional Ethics Committee (AZ D437/09, D451/12, D474/13).

## Author contributions

VK, CK and DK conceived of the research idea and designed the research study. DHM, CK and VK conducted the experiments and acquired data. DHM, VK and MR analysed the data. EE performed the HLA typing for the lymphocyte immunotherapy cohort. VK, CJT and AJB conceived the research programme and provided input into design and the analysis plan. DHM and VK authored the manuscript. All co-authors provided review and revisions to the manuscript and ultimately approved the final version for submission and publication.

## Acknowledgments

This study was supported by the Cambridge NIHR Biomedical Research Centre and the NIHR Blood and Transplant Research Unit in Organ Donation and Transplantation at the University of Cambridge in collaboration with Newcastle University and in partnership with NHS Blood and Transplant (NHSBT). The views expressed are those of the authors and not necessarily those of the NHS, the NIHR, the Department of Health or NHSBT. VK was supported by an Academy of Medical Sciences Grant, an Evelyn Trust Grant and an NIHR Post-Doctoral Fellowship (PDF-2016-09-065). DHM was supported by an RCSEng Research Fellowship. This study was supported by an intramural research grant of Kiel University Medical Faculty (CK) and the Deutsche Forschungsgemeinschaft grant KA 502/18-1 (DK).

